# Stromal localization of inactive CD8^+^ T cells in metastatic mismatch repair deficient colorectal cancer

**DOI:** 10.1101/2023.10.09.561039

**Authors:** Emre Küçükköse, Matthijs J.D. Baars, Mojtaba Amini, Suzanna J. Schraa, Evelien Floor, Guus M. Bol, Inne H.M. Borel Rinkes, Jeanine M.L. Roodhart, Miriam Koopman, Jamila Laoukili, Onno Kranenburg, Yvonne Vercoulen

## Abstract

**Background:** The determinants of metastasis in mismatch repair deficiency with high levels of microsatellite instability (MSI-H) in colorectal cancer (CRC) are poorly understood. Here, we hypothesized that distinct immune and stromal microenvironments in primary tumors may discriminate between non-metastatic MSI-H CRC and metastatic MSI-H CRC.

**Methods:** We profiled 46,727 single cells using high-plex imaging mass cytometry and analyzed both differential cell type abundance, and spatial distribution of stromal and immune cells in primary CRC tumors with or without metastatic capacity. We validated our findings in a second independent cohort using immunohistochemistry.

**Results:** High-plex imaging mass cytometry and hierarchical clustering based on microenvironmental markers separated primary MSI-H CRC tumors with and without metastatic capacity. Primary tumors with metastatic capacity displayed a high stromal content and low influx of CD8^+^ T cells, which expressed significantly lower levels of markers reflecting proliferation (Ki67) and antigen-experience (CD45RO) compared to CD8^+^ T cells in non-metastatic tumors. CD8^+^ T cells showed intra-epithelial localization in non-metastatic tumors, but stromal localization in metastatic tumors, which was validated in a second cohort.

**Conclusion:** We conclude that localization of phenotypically distinct CD8^+^ T cells within stroma may predict metastasis formation in MSI-H CRC.

## Introduction

A deficient DNA mismatch repair system (dMMR) is the underlying cause of a specific subtype of colorectal cancer (CRC). A direct consequence of dMMR is a high level of microsatellite instability (MSI-H). Both can be measured in clinical practice as part of the pathological classification of CRC^1^. In addition, dMMR causes a high mutational load, the generation of tumor-specific immunogenic neo-antigens, and infiltration by T cells. High numbers of tumor-infiltrating T cells are associated with a good prognosis and decreased risk of metastasis^2,3^. Indeed, patients with primary MSI-H CRC have a reduced risk of developing distant metastases^4^. Nevertheless, in a small proportion of patients with MSI-H CRC, distant metastasis formation does occur. These patients account for 3-5% of all cases of metastatic CRC. Importantly, MSI-H status in metastatic CRC is associated with poor prognosis and reduced benefit from chemotherapy^5–7^.

Genetic, histopathological, immunological, and molecular characterization has demonstrated the heterogeneous nature of CRC^8–13^. The association of (some of) these variables with disease progression and therapy response in CRC has promoted precision treatment strategies^14,15^. For instance, MSI-H tumors do not benefit from adjuvant chemotherapy with 5-fluorouacil (5-FU) monotherapy following surgical removal of the primary tumor^16^. Consequently, dMMR tests are now routinely applied in the clinic as a decision tool to select stage II/III CRC patients for adjuvant treatment. Recently, we demonstrated that MMR status in patient-derived CRC epithelial organoid cultures does not correlate with intrinsic tumor cell sensitivity to 5-FU^17^. As organoid cultures lack cells from the tumor microenvironment (TME), we proposed that differences in the TME between MSI-H and microsatellite stable (MSS) tumors may determine benefit from chemotherapy^17^. Indeed, the composition of the TME is now recognized as an important determinant of treatment response in cancer in general^18^.

When compared to MSS tumors, the TME of MSI-H tumors contains high levels of tumor-infiltrating lymphocytes, which are kept in check by expression of immune checkpoints^11,19–22^. Different immune cells (e.g., macrophages, NK-, T- and B cells) in the TME can have pro- or anti-tumorigenic effects. Particularly, reduced infiltration of CD8^+^ cytotoxic T cells into the primary tumor is associated with disease progression and metastasis in CRC (and other cancer types)^2,3^. This observation has formed the basis for developing the ‘Immunoscore’ as a tool to predict CRC progression^2,3^. In addition, a high content of stromal fibroblasts characterizes the metastasis-prone consensus molecular subtype 4 (CMS4) in primary CRC, which predominantly consists of MSS tumors^8,23,24^. To date, prognostic tumor- or TME-centric biomarkers that predict metastasis formation in the MSI-H CRC subtype are lacking and, given the therapy resistance in a substantial proportion of patients with advanced metastatic MSI-H CRC^25^, these are urgently needed to allow for early treatment.

Here, we investigated whether distinct stromal features such as the number, location, and activation status of immune cell subsets and fibroblasts could distinguish between primary MSI-H CRC non-metastatic tumors and tumors with metastatic capacity. To this end, we applied high-plex imaging mass cytometry (IMC), which enables single-cell spatial analysis of multiple distinct cell types in patient-derived tissue samples. We show that metastatic capacity in MSI-H CRC is associated with limited (intra-epithelial) infiltration by CD8^+^ T cells and localization of inactive CD8^+^ T cells within the primary tumor stroma. These findings suggest that a stroma-rich immunosuppressive TME may drive metastasis in MSI-H CRC and may help to devise strategies aimed at early identification and treatment of metastasis-prone MSI-H CRC.

## Results

### TME markers distinguish non-metastatic from metastatic colorectal tumors

We first set out to investigate whether tumor microenvironment features differ between non-metastatic (MSI-H) and metastatic (MSI-H or MSS) primary colorectal tumors. We employed tissue microarrays of patients with non-metastatic MSI-H (*n*=3), metastatic MSI-H (*n*=3) and metastatic MSS (*n*=3) CRC and processed three distinct TMA cores per patient (**Supplementary Table 1**). All cores (*n*=27 Regions of Interest (ROI’s)) were used for multiplex imaging mass cytometry analysis (IMC), using a 35-plex antibody panel (**Supplementary Table 3** ^26^). The panel included markers identifying distinct cell types (i.e. immune cells and fibroblasts), transcription factors, activation/exhaustion markers, and signaling markers (**Fig. 1a, Supplementary Fig. 1**). We used machine learning tools Ilastik and CellProfiler for pixel classification, and tissue region and single cell segmentation^26–29^, and excluded the necrotic areas in ROI’s, which showed non-specific signals with poor signal-noise ratio (**Supplementary Fig. 2a, Supplementary Table 4**). Next, we generated single cell data of all segmented stromal (immune and fibroblast) cells (**Supplementary Fig. 2b**), and normalized signal intensity variability for each marker across different ROIs.

**Fig. 1:**
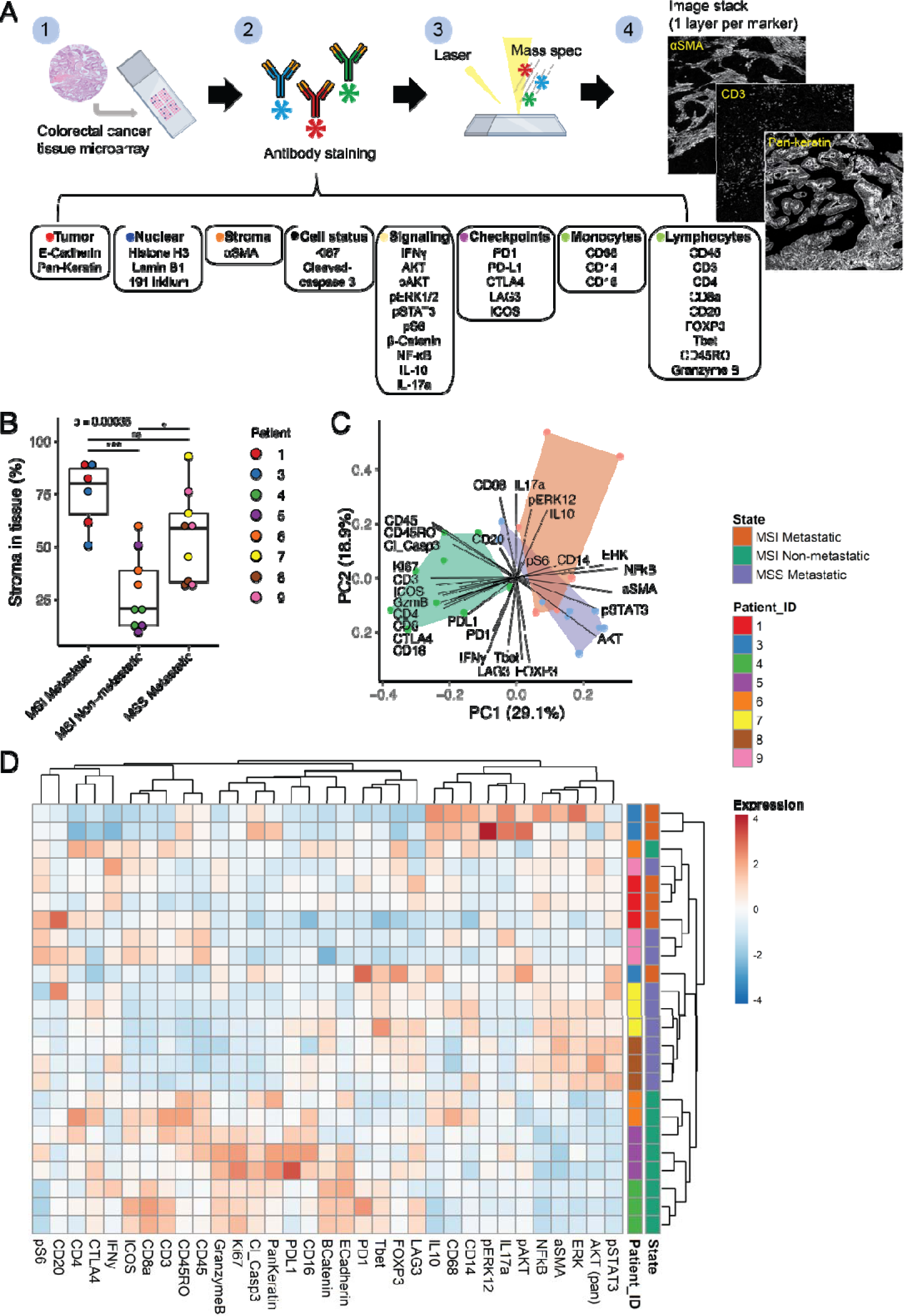
TME markers distinguish non-metastatic from metastatic colorectal tumors. (**a**) Illustration of the data acquisition workflow used for Imaging mass cytometry (IMC): (1) One 4 µm-thick section of a TMA composed of 2 mm^2^ CRC cores (2) was stained with a cocktail of 35 antibodies labeled with unique metal isotopes. (3) Samples were ablated with a high-energy laser in a rasterized pattern. The resulting plumes were ionized and analyzed by a mass cytometer, which returned the metal isotope composition per pixel. (4) Each antibody resulted in a single image per sample, and together constructed a multi-channel image stack. (**b**) Stromal content was quantified by analyzing α-Smooth muscle actin (aSMA) expression. Boxplot represents the percentage of aSMA^+^ stroma (fibroblasts) in tissue per group, individual patients are color-coded. (**c**) Principal component analysis using the median intensity of each marker per image region as input variables. Each data point is a given ROI, color-coded by tumor state. Black lines depict the contribution of individual markers to the first two principal components (PC1 and PC2) of the analysis. (**d**) Hierarchical clustering analysis of ROIs according to median marker intensity of segmented stromal/immune cells. Tumor state and patient ID are indicated using a color code on the right side.

Analysis of segmented epithelial and stroma compartments showed that the stroma content (percentage of stroma in the tissue) was significantly higher in MSS/MSI-H metastatic tumors compared to MSI-H non-metastatic tumors (**Fig. 1b**). Principal component analysis showed separate clustering of metastatic and non-metastatic tumors based on differential marker expression (**Fig. 1c**). Hierarchical clustering revealed a separate cluster containing most non-metastatic tumors (**Fig. 1d**). Both analyses demonstrate that, in comparison to metastatic tumors (MSS and MSI-H), non-metastatic MSI-H tumors display high expression of markers for proliferation (Ki67) and cell death (cleaved caspase 3), and of markers identifying immune cells (CD45, Granzyme B), including T cells (CD3, CD8, CD45RO, ICOS, CTLA-4) (**Fig. 1c, d** and **Supplementary Fig. 3a).** Moreover, MSI-H non-metastatic tumors show low expression of the stromal marker aSMA, and markers associated with inflammation (phosphorylated STAT3 (P-STAT3), NFkB) (**Fig. 1d**).

### Metastatic MSI-H tumors contain low numbers of cytotoxic T cells

To gain further insight into the differences in TME composition between metastatic and non-metastatic tumors, we performed unsupervised clustering by Phenograph^30^, and manually annotated clusters of cell types based on marker expression (**Supplementary Fig. 3b)**. We identified 20 clusters and 12 distinct cell types (**Fig. 2a**). Uniform Manifold Approximation and Projection (UMAP) analysis on the expression of lineage markers per cell showed a clear separation of the identified cell types (**Fig. 2b**). Hierarchical clustering analyses of cell type abundance revealed strong predominant ordering according to disease state and showed two major clusters (**Fig. 2c**). The first cluster mainly consisted of MSI non-metastatic tumors and displayed elevated proportions of T cell populations (cytotoxic, helper and regulatory), while the second cluster mainly contained MSI-H/MSS metastatic tumors and displayed higher abundance of ‘Fibroblast’, ‘Fibroblast & Monocyte’, ‘Neutrophil & NK-cell’ populations (**Fig. 2c**). Indeed, in line with the higher stroma content observed in metastatic tumors (**Fig. 1b**), MSS metastatic tumors showed significantly elevated numbers of ‘Fibroblast’ and ‘Fibroblast and Monocyte’ populations compared to MSI-H non-metastatic tumors (**Fig. 2d**). MSI-H non-metastatic tumors contained a significantly higher number of ‘Cytotoxic T cell’, ‘Regulatory T cell’, and ‘Macrophage & Monocyte’ when compared to MSI-H metastatic tumors (**Fig. 2d**). ‘B cells’, ‘T helper cells’, and other cell types were present in equal numbers in MSI-H non-metastatic and MSI-H metastatic tumors (**Fig. 2d**).

**Fig. 2:**
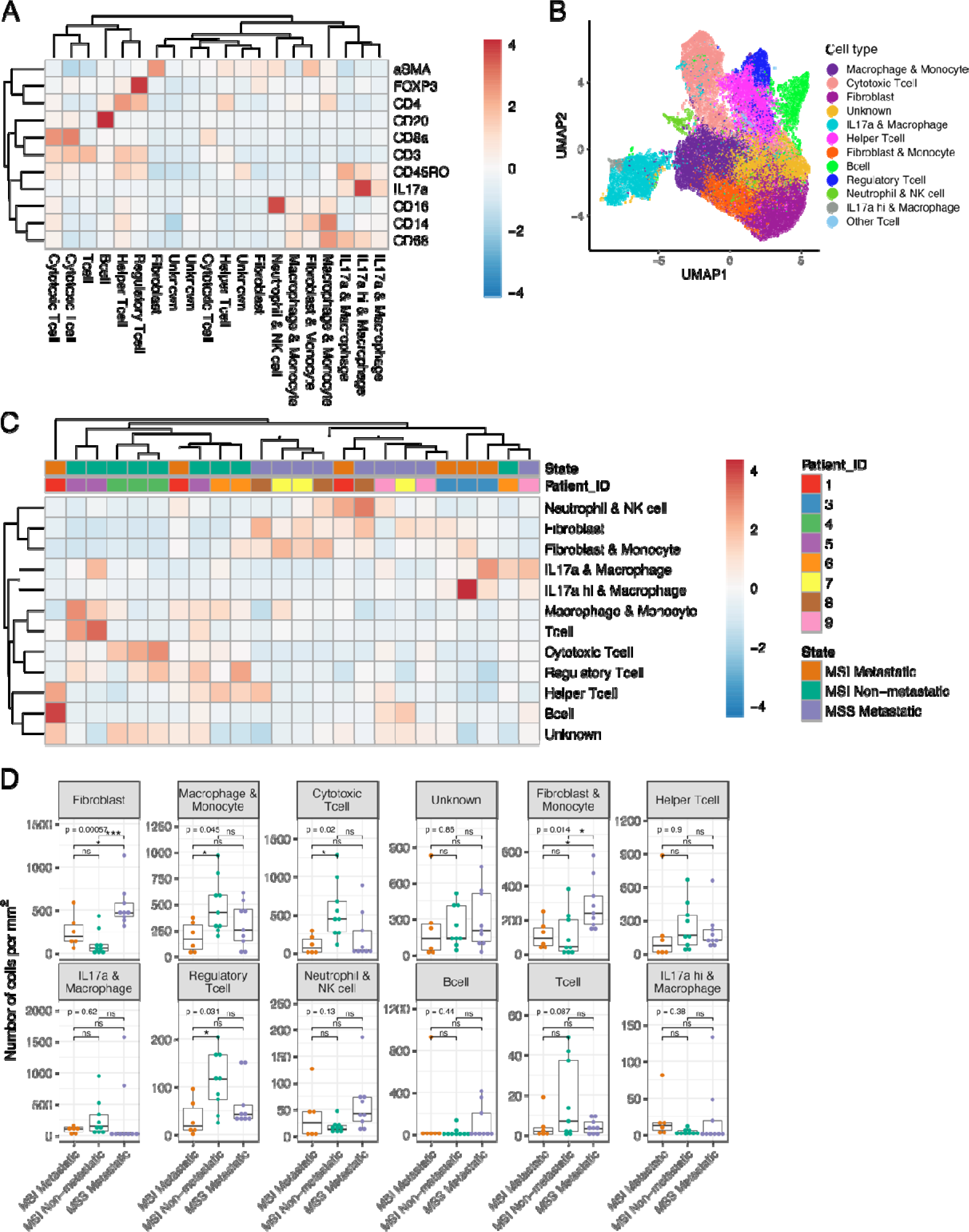
Cell-type annotation reveals differential microenvironment composition between metastatic and non-metastatic primary CRC tumors. (**a**) Hierarchical cluster analysis heatmap of lineage marker expression. The heatmap shows the expression levels of 11 markers used for defining cell clusters. (**b**) Uniform Manifold Approximation and Projection (UMAP) of single cells (*N*=46,727) of all ROIs combined using lineage marker intensities, color-coded by annotated cell lineages. (**c**) Heatmap showing hierarchical clustering on cell-type abundance for all cell lineages and individual ROIs. (**d**) Boxplot showing the median of number of cells per mm^2^ stratified for annotated cell-types, across disease groups

### Metastatic MSI-H tumors lack proliferating antigen-experienced CD8**^+^** T cells

Next, we investigated the relative proportions of specific T cell sub-populations (CD4^+^, CD8^+^ and FOXP3^+^) in the tumors. This analysis showed that the CD8^+^ T cell population was more abundant in MSI-H non-metastatic tumors compared to MSS metastatic tumors (**Fig. 3a**). *Vice versa*, the proportion of CD4^+^ T cells was higher in MSS metastatic tumors compared to MSI-H non-metastatic tumors. The proportion of regulatory T cells (Tregs; FOXP3^+^) was similar in all tumors (**Fig. 2d**, **Fig. 3a**).

**Fig. 3:**
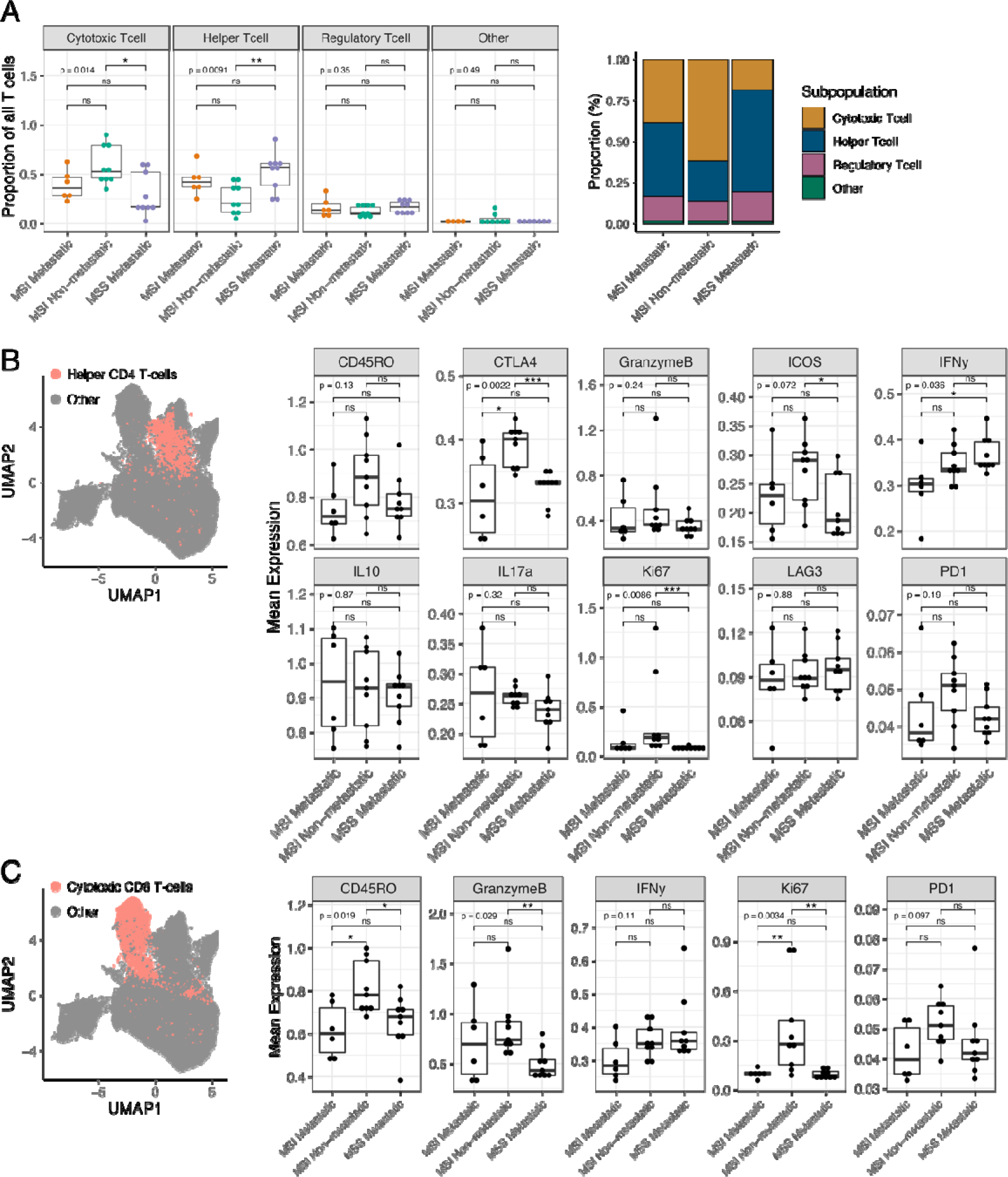
Metastatic MSI-H tumors lack proliferating antigen-experienced CD8^+^ T cell s. **(a)** Boxplot and stacked bar plot represent the proportions of cytotoxic-, helper-, regulatory- or other T cell of all T cells. (**b**) UMAP of all stromal and immune cells, highlighting the helper T cells as red dots. Boxplots represent the median expression per ROI of given markers for helper T cells. (**c**) UMAP of all stromal and immune cells, cytotoxic T cells are indicated as red dots. Boxplots represent the median expression of given markers per ROI for cytotoxic T cells.

Analysis of immune checkpoint expression showed that CTLA-4 was significantly higher in helper T cells in MSI-H non-metastatic tumors compared to MSI-H/MSS metastatic tumors (**Fig. 3b**). Other proteins expressed by T helper cells (CD45RO, Granzyme B, IFNy, IL10, IL17a, Ki67, LAG3, PD1) were not significantly different between the three groups. Tregs showed no changes in expression of immune checkpoints and displayed significantly increased proliferation (Ki67) in non-metastatic MSI-H CRC compared to the metastatic tumors (**Supplementary Fig. 4**). In cytotoxic T cells, expression of the effector molecule Granzyme B was elevated in MSI-H non-metastatic tumors compared to MSS metastatic tumors. Moreover, CD45RO and Ki67 expression in cytotoxic T cells were increased in MSI-H non-metastatic tumors compared to both MSI-H and MSS metastatic tumors (**Fig. 3c**).

### Stromal, but not intra-epithelial, localization of cytotoxic T cells in metastatic MSI-H tumors

We observed that the immune cells in MSI-H and MSS metastatic tumors appeared to be excluded from the tumor compartment compared to non-metastatic tumors. To quantify this, we determined the distance towards the epithelial tumor cells of all distinct cell populations in the TME (fibroblasts, B cells, T cells, monocytes, neutrophils, NK cells). To this end, we constructed a spatial graph in which individual cell-centroids are represented as dots and segmented tumor regions are converted into polygons (**Fig. 4a**, **Supplementary Fig. 4**). The minimal distance between each given single cell and the tumor outline was then determined for each cell type. The distance between the tumor compartment and ‘Fibroblasts’, ‘B cells’, ‘Fibroblast & Monocyte’, and ‘Neutrophils & NK cell’ were similar between groups. However, T cell populations, including cytotoxic T cells, localized very close to the tumor cells in MSI-H non-metastatic tumors (median 2 µm distance) but significantly further away from the tumor cells in MSI-H/MSS metastatic tumors (median distance 21 and 22 µm; *p*=2e-16) (**Fig. 4b, c, Supplementary Fig. 6**). Similarly, macrophage-containing cell populations were localized near the tumor cells in MSI-H non-metastatic tumors, but further away from the tumor cells in metastatic tumors (**Fig. 4b**).

**Fig. 4:**
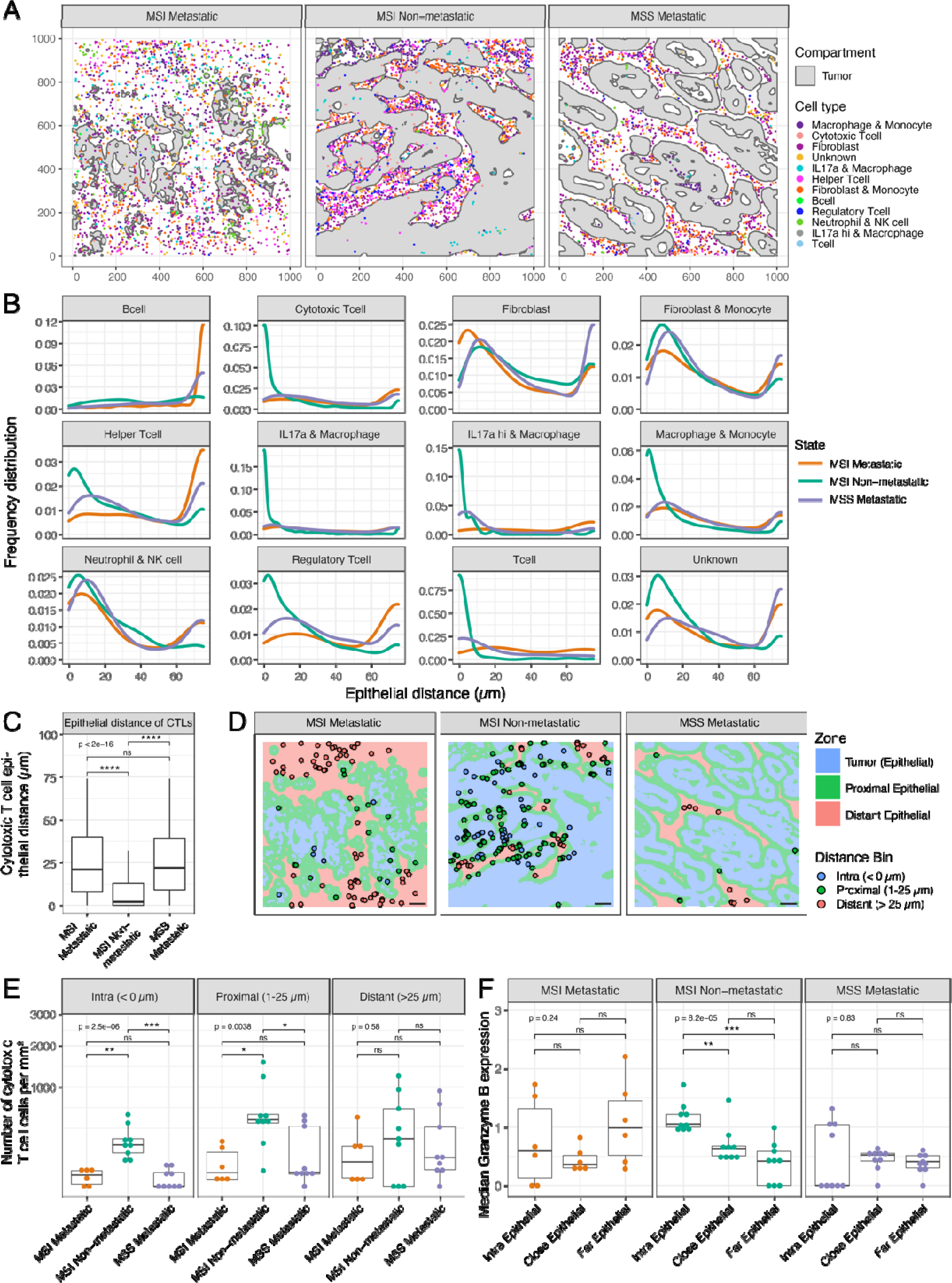
Spatial analysis reveals distinct localization of cell-types in the tumor microenvironment. (**a**) Spatial representation of representative ROIs per disease group with segmented tumor regions displayed as grey polygons and the identified cells are shown as dots, colored by cell-type. (**b**) Tumor distance plots demonstrating the relative frequency distribution of given cell populations. (**c**) Boxplot represents the median epithelial distance of cytotoxic T cells per group. (**d**) Spatial representation of cytotoxic T cell localization relative to the tumor, color-coded by distance bin: ‘intra’ (blue), ‘proximal’ (1 – 25 µm; green) and ‘distant’ (>25 µm; red). Scale bar indicates 100 µm. (**e**) Boxplots represent the number of cytotoxic T cells per mm^2^ per distance bin and group (median per ROI). (**f**) Boxplots represent the median Granzyme B expression in cytotoxic T cells per distance bin and group (median per ROI).

To assess cytotoxic T cell numbers in relation to their distance from the tumor compartment, we divided the cytotoxic T cell population in three distinct distance-based bins, “intra-epithelial” (≤ 0 µm), “proximal” (1 – 25 µm), and “distant” (> 25 µm) (**Fig. 4d**). The accumulation of T cells in the ‘intra-epithelial’ and ‘proximal’ bins was significantly higher in MSI-H non-metastatic tumors when compared to MSI-H/MSS metastatic tumors, while ‘distant’ T cells were similar among the groups (**Fig. 4e**). The levels of the ‘intra-epithelial’ tumor infiltrating cytotoxic T cells clearly differentiated non-metastatic from metastatic tumors, and this difference appeared more robust than the difference in total T cell numbers (**Fig. 3a**). Moreover, intra-epithelial T cells expressed significantly higher levels of Granzyme B when compared to T cells located further away from the tumor cells in MSI-H non-metastatic tumors. This was not observed in MSI-H or MSS metastatic tumors (**Fig. 4f**).

To validate the relevance of the robust difference in cytotoxic T cell localization in the TMA samples, we validated our findings in a second independent cohort of patients with MSI-H metastatic disease (n=10) or MSI-H non-metastatic disease (n=11), using large tumor sections (**Fig. 5a**). We performed immunohistochemistry staining for the detection of epithelial tumor cells (E-cadherin), stromal fibroblasts, (αSMA) and cytotoxic T cells (CD8a) (**Fig. 5b**). Cytotoxic T cells in non-metastatic MSI-H tumors localized significantly (p=0.00012) closer to tumor cells than those in metastatic MSI-H tumors (**Fig. 5c, d)**. Stromal content was significantly (p=0.029) higher in MSI-H metastatic tumors than in non– metastatic tumors (**Fig. 5e**).

**Fig. 5:**
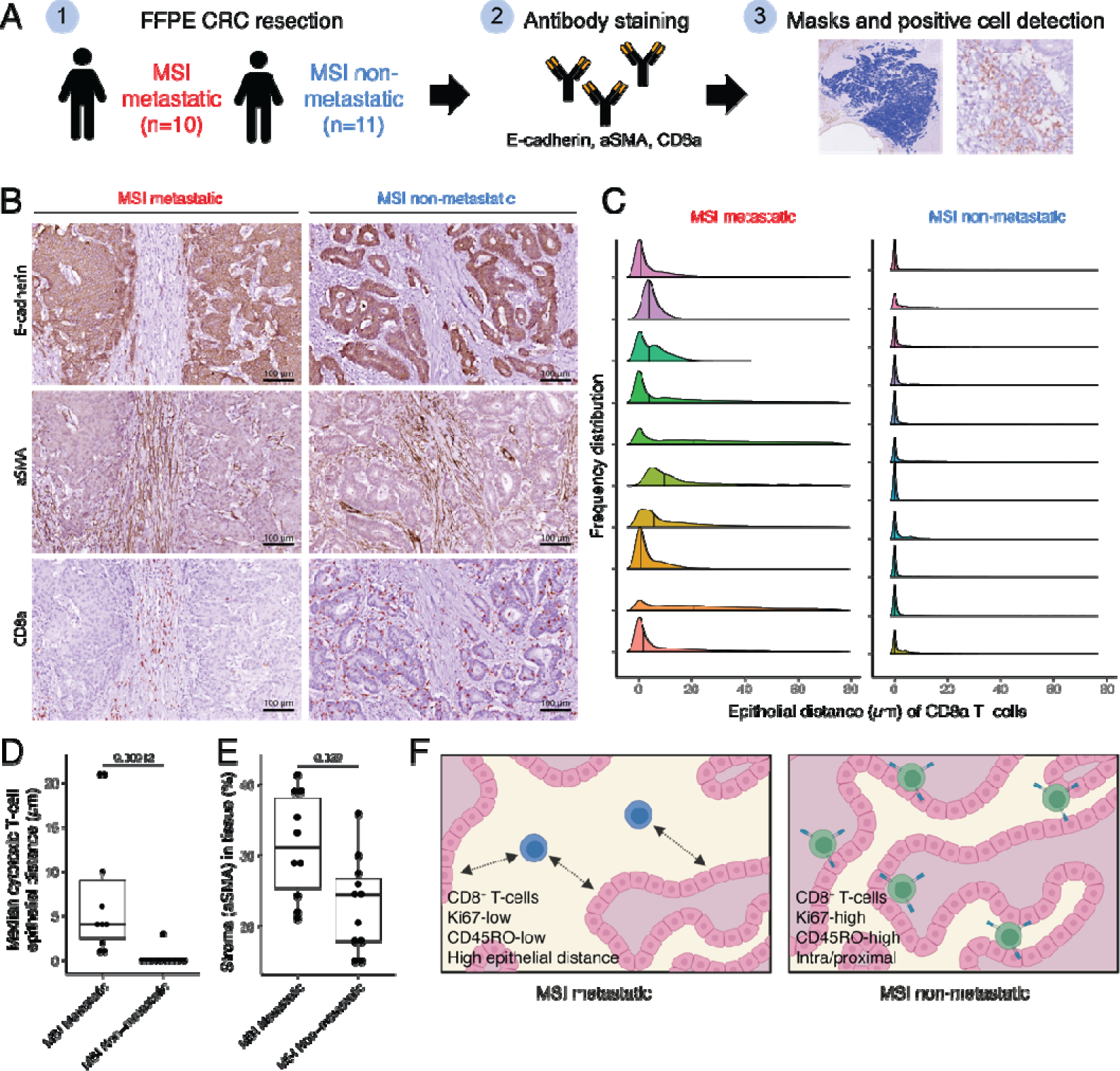
Cytotoxic T cells reside in high stromal content in metastatic MSI-H tumors. (**a**) Illustration of the data acquisition workflow used for the IHC validation cohort: (1) FFPE sections of resected primary CRC with (*n*=10) or without (*n*=11) metastatic disease were (2) stained for E-cadherin, aSMA and CD8a using IHC. (3) Tumor segmentation was performed, and masks were generated using Qupath^43^. Individual cytotoxic T cells were identified. (**b**) Representative IHC images of E-cadherin, aSMA, and CD8a. Scale bar is 100 µm. (**c**) Tumor distance plots demonstrating the frequency distribution of cytotoxic T cells per tumor and group. (**d**) Boxplot representing the median epithelial distance of cytotoxic T cells per ROI, shown per group. (**e**) Boxplot representing the percentage of stroma per ROI and group. (**f**) Schematic model of CD8 T cell phenotype and localization in the CRC microenvironment of metastatic and non-metastatic MSI-H tumors.

Together, our findings support a model in which metastatic primary MSI-H tumors characterized by extensive formation of fibroblast-rich stroma in which cytotoxic T cells are residing and remain inactive, while in non-metastatic MSI-H tumors the cytotoxic T cells localize intra-epithelially, close to tumor cells, proliferate, and have experienced tumor antigen (**Fig. 5f**).

## Discussion

In this study, we profiled 46,727 single cells using multiplex imaging mass cytometry and analyzed both differential cell type abundance, and spatial distribution of stromal and immune cells in metastatic and non-metastatic primary colorectal tumors. We demonstrate that MSI-H tumors with metastatic capacity are characterized by extensive formation of fibroblast-rich stroma containing inactive cytotoxic T cells which fail to invade the tumor cell compartment. By contrast, non-metastatic MSI-H tumors were characterized by infiltrating and actively proliferating antigen-experienced cytotoxic T cells (CD8^+^CD45RO^+^Ki67^+^), which produced high levels of Granzyme B, indicating cytotoxic activity. Moreover, these tumors displayed a higher number of proliferating Tregs, and helper T cells with elevated expression of immune checkpoint molecules (CTLA-4). These results show that both anti-tumor and immune-suppressive responses occur in the microenvironment of non-metastatic MSI-H tumors, with a predominantly cytotoxic T cell response.

T cell infiltration and activation is associated with a better prognosis in CRC^2,3,31^, and our findings confirm the prognostic value of cytotoxic T cell infiltration for metastatic MSI-H CRC. The numbers of cytotoxic T cells detected might be less reliable due to location-dependent bias and sample variation, in particular when analyzing relatively small tumor regions. Here, we show that intra-tumoral localization of T cells is a more robust method. Our results are in line with a correlation between intra-tumoral localization of cytotoxic T cells with a memory phenotype, and survival demonstrated in breast cancer^32^, and confirm the value of spatial analysis of the tumor microenvironment^33^. Tumors containing high numbers of fibroblasts often have low levels of lymphocytes^34,35^. A recent study demonstrated that specific subsets of cancer-associated fibroblasts with each unique molecular programs and organization patterns, associate with T cell exclusion in human lung cancers^36^. Increased stroma is generally associated with worse overall survival and recurrence in CRC patients after surgery^23,24,37^. Stromal localization of cytotoxic T cells has thus far not been associated with the risk of metastasis formation in CRC patients. Our study suggests that tumor stroma may contribute to metastasis by trapping cytotoxic T cells and preventing T cell expansion and activation.

Immune Checkpoint Blockade (ICB) therapy has led to durable responses and significantly improved overall survival, and now replaces first-line chemotherapy in metastatic MSI-H CRC patients^38–40^. Furthermore, neoadjuvant PD-1 and CTLA-4 ICB therapy in primary colon cancers (stage I-III) showed that MSI-H tumors responded to ICB (*n*=20), and the presence of CD8^+^PD1^+^ T cells predicted ICB response^41^. Here, our spatial analysis revealed that primary MSI-H tumors with metastatic capacity were largely devoid of intra-epithelial cytotoxic T cells. The success of ICB in relation to T cell intra-epithelial versus stromal localization in the tumor remains to be investigated in the future.

In conclusion, our study reveals that primary MSI-H tumors that gave rise to metastases are characterized by low numbers of inactive cytotoxic T cells localized in the stroma, in both early (I-III) and late tumor stages (IV). We propose that combined histopathological evaluation of stroma content, cytotoxic T cell activation status and spatial localization, may have value in predicting metastasis formation in early-stage MSI-H CRC. Such prognostic value should now first be determined in larger patient cohorts. The next step would be to develop a robust metastasis-prediction test for MSI-H CRC which may have significant clinical utility in terms of patient selection for intensified follow-up and/or (neo) adjuvant (immuno)therapy.

## Methods

### Human tissues

We generated a collection of primary tumor tissues derived from metastatic MSI-H-CRC (*n*=3; CAIRO3 study^42^), non-metastatic MSI-H CRC (*n*=3; UMCU biobank) and metastatic MSS CRC (*n*=3; CAIRO3 study^42^) (**Supplementary Table 1**). Patients did not receive systemic therapy before resection. The collection and processing of human tissue specimens used for validation of our findings (**Supplementary Table 2**) was performed in accordance with the Declaration of Helsinki and approved by the Medical Research Ethics Committee of Utrecht, the Netherlands (METC 20-102) and Dutch Nationwide Pathology Databank (PALGA 2020-53). This tissue is classified as “residual material” and the collection and processing of this biological material is in accordance with the “no objection” procedure defining the release of anonymized residual material without broad consent under strict conditions and approval under afore-mentioned Research Ethics Committee. One patient (out of 21) received capecitabine + oxaliplatin treatment before resection.

### Antibodies and reagents

For a comprehensive list of antibodies, compounds, and kits, see **Supplementary Table 3**^26^. **Immunohistochemistry** Tumor sections of 4 µm thickness were deparaffinized and rehydrated. Endogenous peroxidase activity was blocked with 1.5% hydrogen peroxide for 10 min. Heat-induced antigen retrieval was carried out using citrate buffer pH6.0 for 20 min, followed by cooling of tissue sections for 20 min. Sections were incubated overnight at 4°C with primary antibodies (**Supplementary Table 3**) diluted in PBS with 0.1% Sodium azide and 3% BSA. The next day, sections were washed three times with 0.05% Tween-PBS solution for 5 min, followed by a 1 hour incubation with a HRP-conjugated secondary antibody. Subsequently, sections were washed three times with 1× PBS for 5 min and developed with 3,3’-Diaminobenzidine (DAB) chromogen for 10 min at room temperature. Sections counterstained with hematoxylin, followed by dehydration and mounting.

Stained tissue sections were scanned using NanoZoomerXR whole slide scanner (Hamamatsu) at 40X magnification, with a resolution of 0.25 μm/pixel. Quantification of the staining was performed using QuPath software^43^. Targets of interest were quantified using QuPath’s Trained Pixel classification command that allows an automated recognition of background, tissue (haematoxylin) and DAB staining areas. The percentage of target in regions of interest (e.g. αSMA for stroma content) was then calculated by using the following formula: DAB area / (DAB + tissue – background area) * 100%.

### Imaging mass cytometry

Imaging mass cytometry was performed exactly as described in the MATISSE protocol^26,29^.

### Single cell segmentation

Segmentation of single cells was performed on IMC data in a two-step process as described before^26,29^, with the following deviations: First, a pixel classification workflow was used to identify nuclei and membranes of stromal cells in all images using IMC data. The resulting probability maps were used to perform single-cell segmentation. Importantly, individual tumor cells were neglected in these steps. The following channels were used for nucleus and membrane pixel identification: Histone H3, DNA-lr193 and CD3, CD4 CD8a, CD14, CD16, CD20, CD45, CD45RO, CD68.

### Tissue compartment segmentation

Tissue type training for identification of tumor, stroma, and background regions was performed in Ilastik with a pixel classification workflow using the following IMC channels: Histone H3, DNA-Ir193, E-Cadherin, Pan Keratin, and Beta-catenin. The following features were selected: Gaussian Smoothing, Laplacian of Gaussian, Gaussian Gradient Magnitude, Differenced of Gaussians, Structure Tensor Eigenvalues, Hessian of Gaussian Eigenvalues (σ3.5, 5, 10).

### Single cell data generation

Single cell data was generated in R (v4.0) by extracting pixel intensities from unscaled 32-bit images for all channels for all cells represented in the segmentation maps, as described before^26,29^.

### Data curation

Necrotic tissue regions were manually excluded from analysis based on IMC and toluidine blue images. High-intensity noise artefacts in IMC images were corrected by excluding single-cell events overlapping with these regions. The artefacts were identified with a pixel classification workflow in Ilastik using all acquired channels. The following feature was selected: Gaussian Smoothing (σ0.3, 1.0). Single-cell events with a mean probability above 0.1 for predicated noise were excluded. Samples from patient 2 were excluded from analysis due to bad tissue quality resulting in poor signal-noise ratio in staining.

### Signal normalization

Intensities were log1p scaled. Next, signal levels for individual image regions were normalized using the median of scaled intensities of the following channels: AKT (pan), CD20, CD45, CD45RO, Cl_Casp3, CTLA-4, ERK (pan), FOXP3, IFNγ, LAG3, NFkB, pAKT, pSTAT3, Tbet.

### Cell type annotation

Single-cell events were clustered using Rphenograph (*k*=20) on normalized and rescaled (R scale function) intensity values using the following lineage markers: aSMA, CD3, CD4, CD8a, CD14, CD16, CD20, CD45RO, CD68, FOXP3, IL17a. Resulting clusters were manually assigned to cell types based on marker expression patterns.

### Expression analysis

Unless otherwise specified, expression values were calculated as median expression of all cells per image. Heatmaps were generated using the pheatmap (RRID: SCR_016418) package in R.

### Spatial analysis

For IMC data, segmented tumor regions were converted to polygons in R using the sf package^44^. Buffer regions around tumor regions were generated using the st_buffer function. Distance to tumor regions are centroid based. In frequency distribution graphs distance is clipped to 75μm.

For IHC data, tumor and stroma regions per image are segmented using a pixel classification workflow in QuPath on CD8a-stained sections. Segmentation was optimized on a single-sample level and verified using a serial E-Cadherin stained section. CD8a cytotoxic T cells were identified using a positive cell detection workflow in QuPath. Single-cell events were converted to centroids, and closest distance to tumor borders was calculated. Events with distances above 75μm were discarded from further analysis.

### Statistics

For cellular frequency analysis we used one-way ANOVA and Tukey HSD post-hoc test. For expression analysis, we used Kruskal-Wallis and Dunn post-hoc test with Bonferroni correction.

## Data availability

All data will be made publicly available in Zenodo repository (https://doi.org/10.5281/zenodo.7692926)

## Code availability

All code will be made publicly available in Github (https://github.com/VercoulenLab).

---

**Supplementary Fig. 1:**
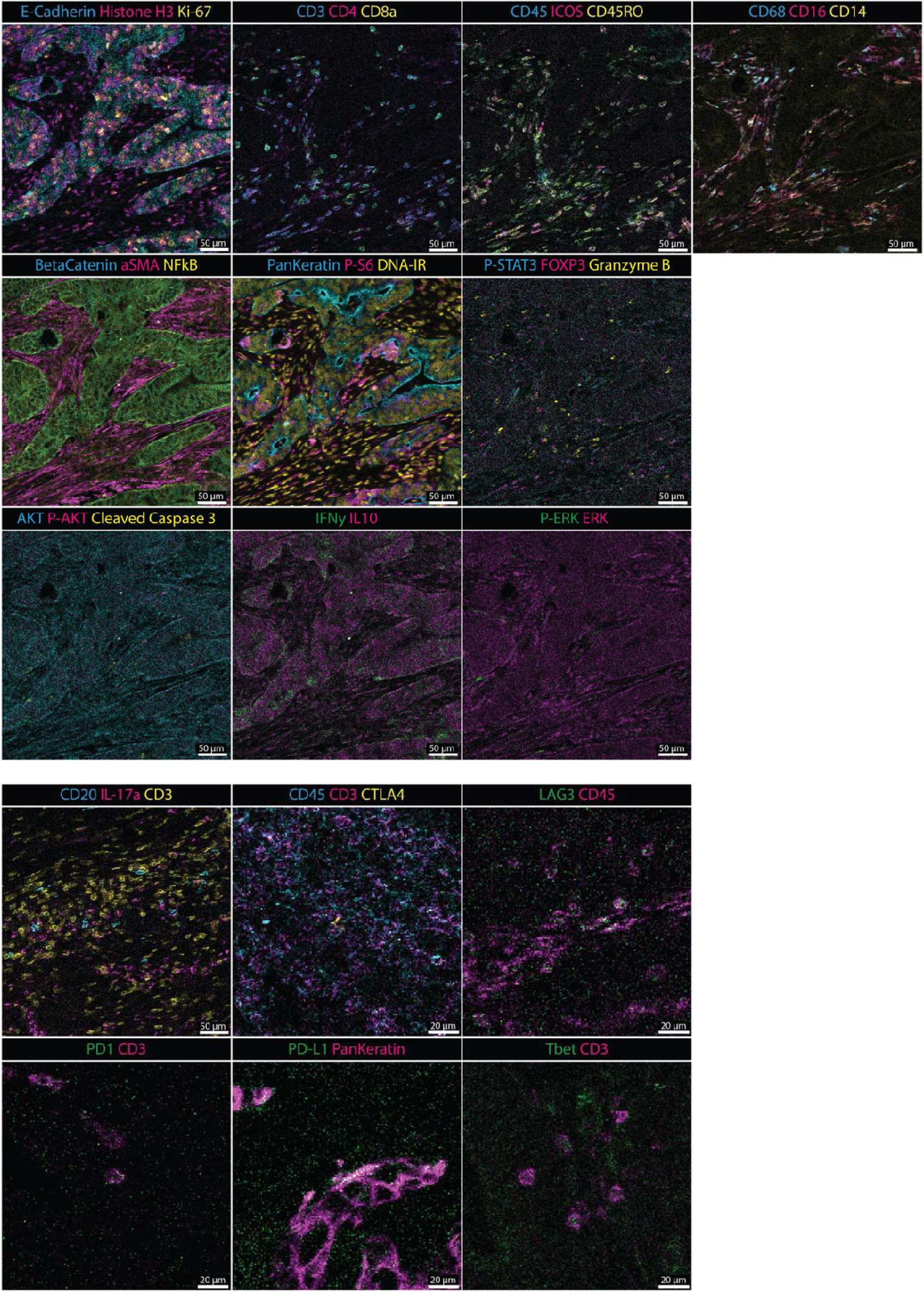
Representative images demonstrating all markers acquired in the multiplex imaging mass cytometry. All 35 antibody-stainings used for IMC are shown. Scale bar sizes are indicated

**Supplementary Fig. 2:**
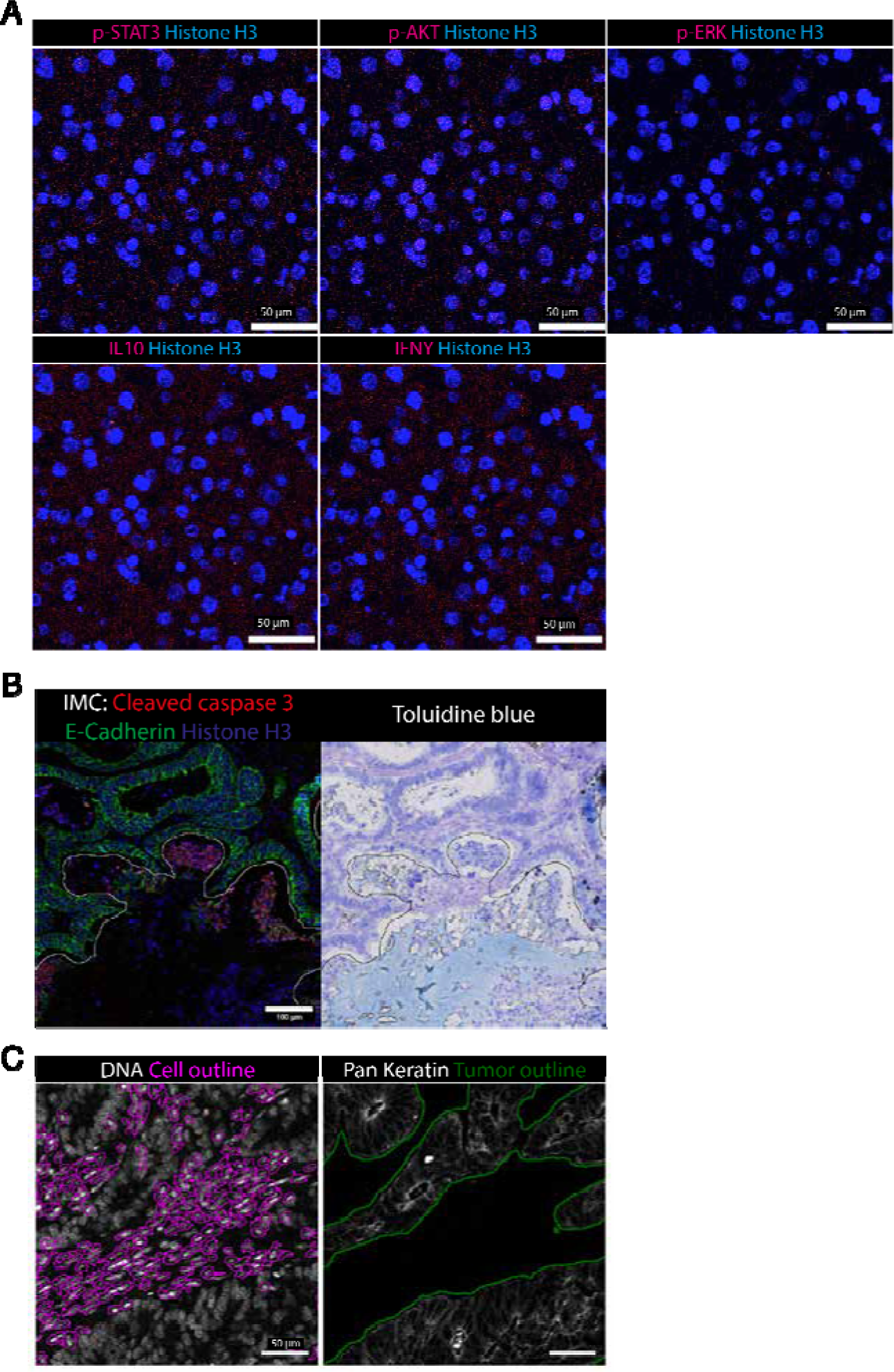
Negative controls, exclusion of necrotic regions, and single-cell and tumor segmentation in IMC data. (**a**) Negative controls for p-STAT3, p-AKT, p-ERK, IL1- and IFNy stainings in IMC. Scale bar is 50 µm. (**b**) Representative ROI demonstrating necrotic area, left: IMC staining data, necrotic region outlined in yellow. Right: Slide-scan image of same section, necrotic region outlined in black (see also **Supplementary Table 4**). Scale bar is 100 μm. (**c**) Representative image showing the predicted outlines for single-cell (magenta) and tumor (green) segmentation.

**Supplementary Fig. 3:**
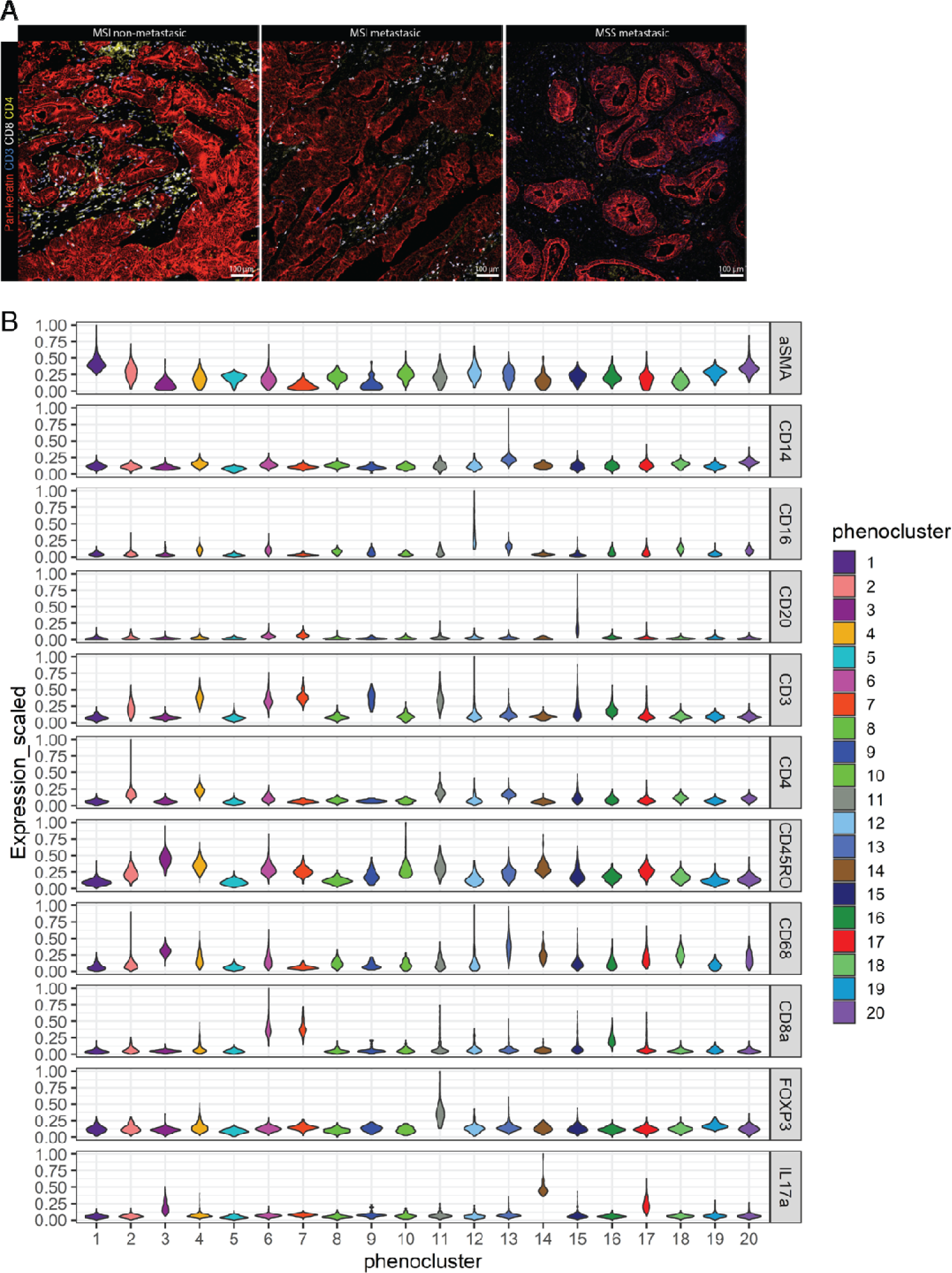
Cell-type annotation by clustering single-cells on lineage marker expression. **(a)** Representative IMC images showing pan-keratin (red), CD3 (blue), CD8 (white) and CD4 (yellow) per group. Scale bar is 100 µm. (**b**) Violin plot representing scaled expression of lineage markers per generated Rphenograph cluster.

**Supplementary Fig. 4:**
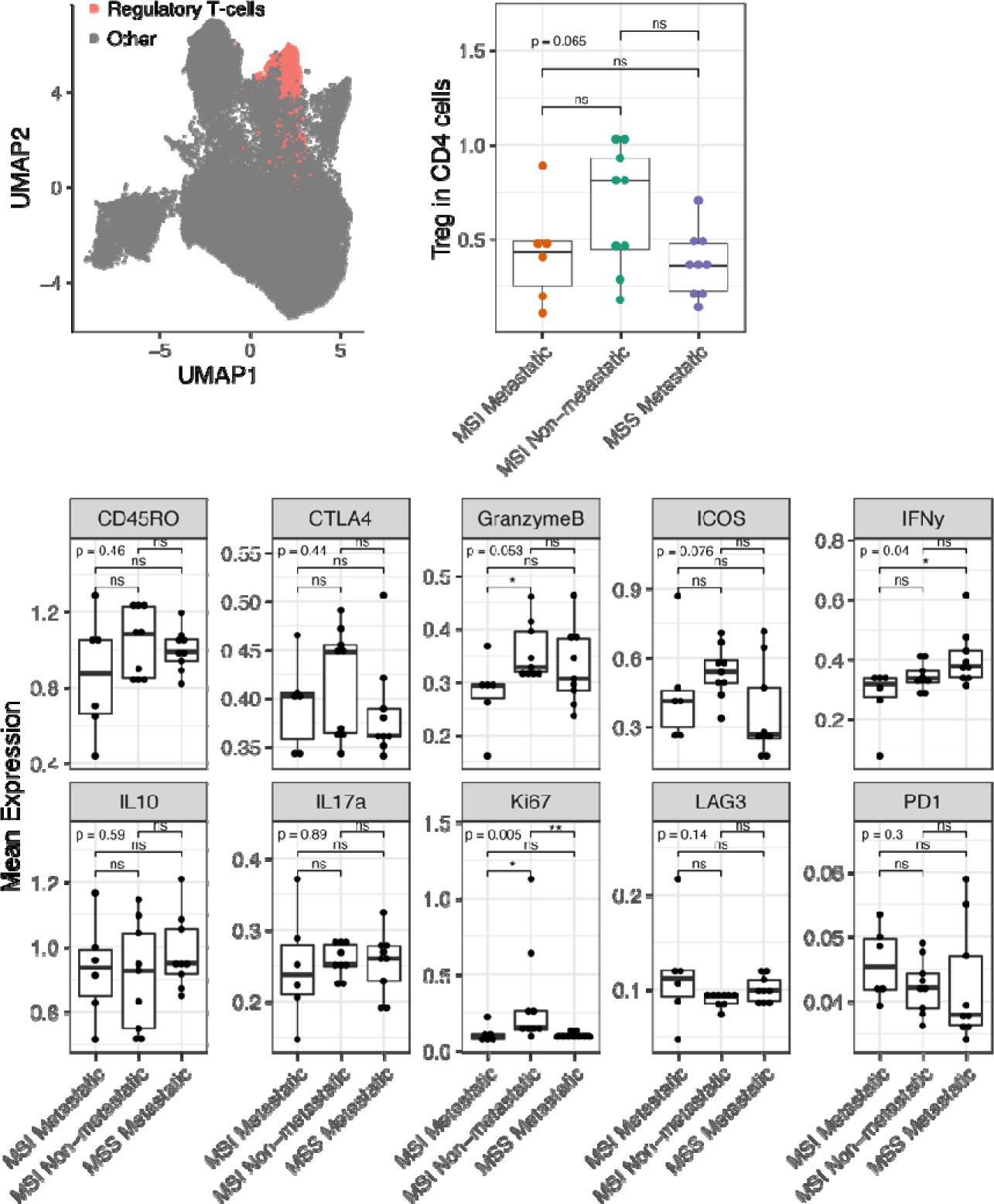
Increased proliferating regulatory T cells in non-metastatic MSI-H tumors compared to metastatic tumors. UMAP of all stromal and immune cells, highlighting the regulatory T cells as red dots. Boxplots represent the median expression per ROI of given markers for regulatory T cells.

**Supplementary Fig. 5:**
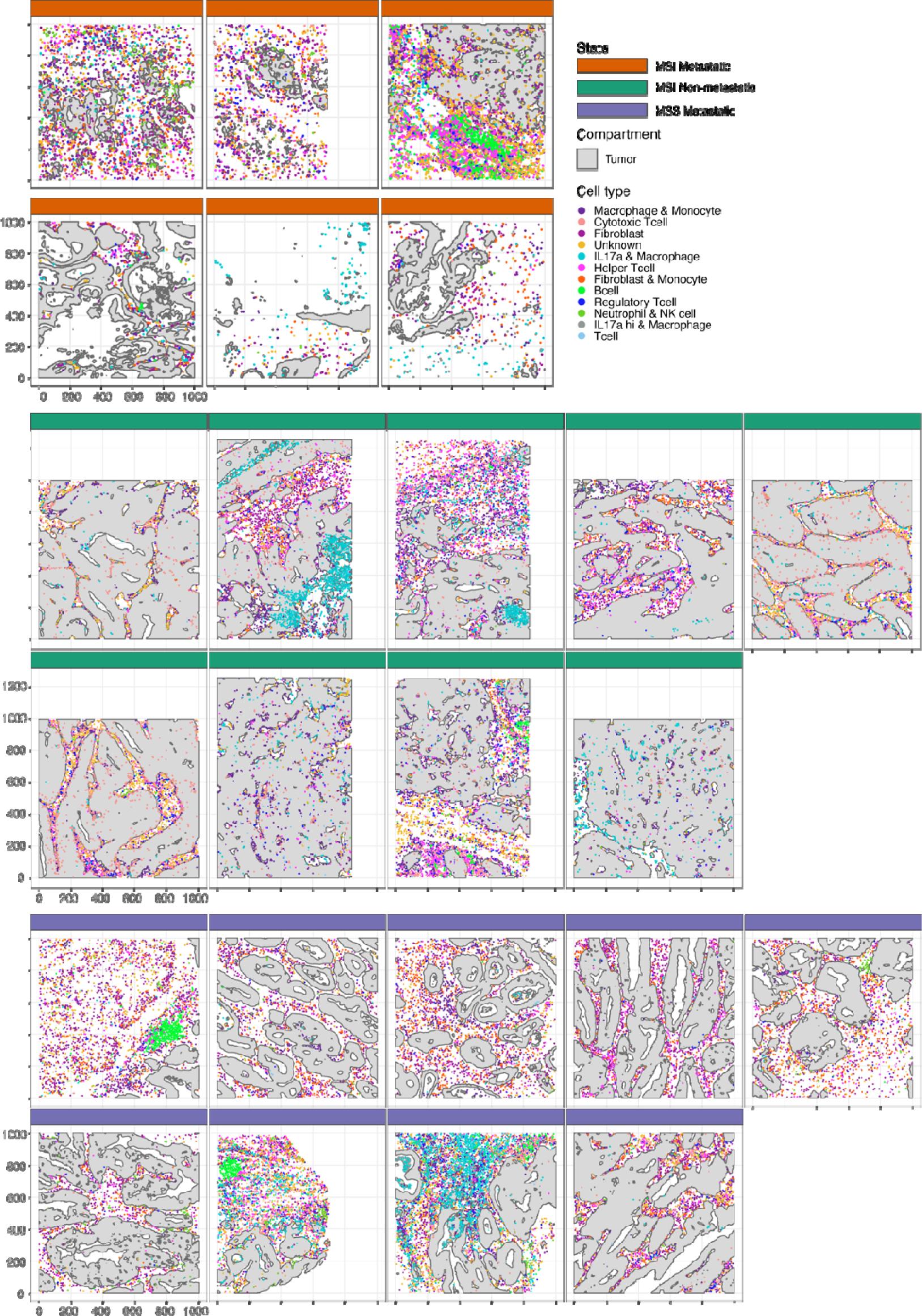
Spatial immune landscape of CRC microenvironment for all samples. Spatial representation of segmented tumor regions shown as grey polygons, and single-cell events shown as dots, colored by the 12 identified cell populations. All ROIs are included.

**Supplementary Fig. 6:**
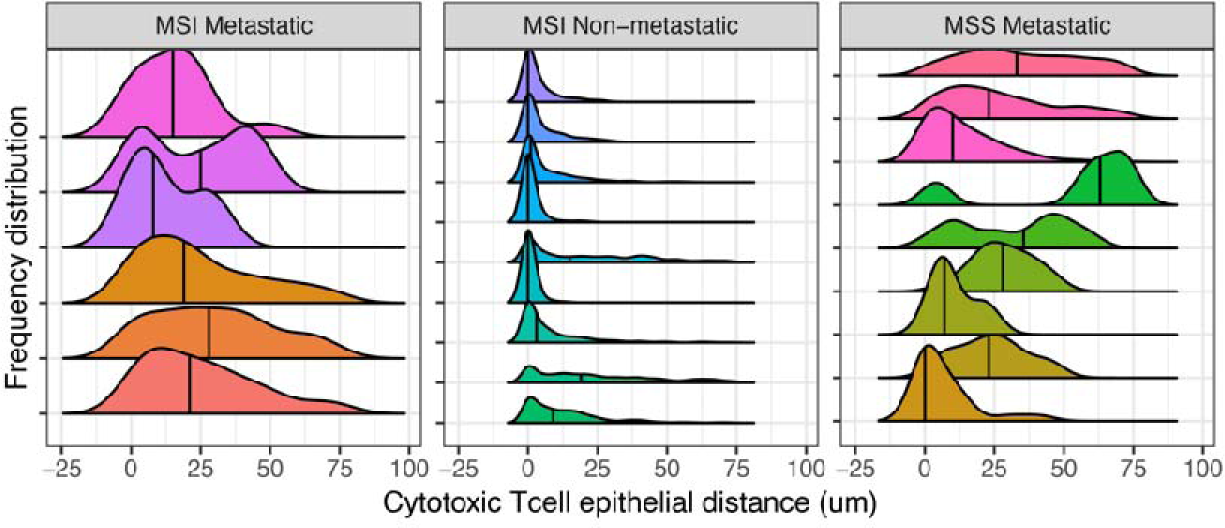
Relative tumor distance of cytotoxic T cells in individual IMC ROIs. Frequency distribution of cytotoxic T cell tumor distance for all CyTOF IMC ROIs, per group.

## ACKNOWLEDGEMENTS

The authors thank all patients involved in the current study.

## AUTHOR CONTRIBUTIONS

Study design: EK, MJDB, OK, YV. Experimental work: EK, MJDB, MA, EF. Resources: SS, GB, JMLR, MK. Data interpretation: EK, MJDB, IHMBR, JL, OK, YV. Study supervision: OK, YV. Writing manuscript: EK, MJDB, OK, YV.

## COMPETING INTERESTS

The authors declare no conflicts of interest.

## FUNDING

This work was supported by the Dutch Cancer Society (KWF/Alpe d’HuZes #UU-10660), and the Dutch Scientific Organization (cancergenomicscenter.nl, NWO Gravitation 024.001.028). Funders had no role in study design, data acquisition, analysis, and interpretation.

## Notes

### Competing Interest Statement

The authors have declared no competing interest.

